# Butyrate synergizes with glucose to promote anaerobic growth of *Staphylococcus aureus* via anaplerotic metabolism and alanine synthesis

**DOI:** 10.64898/2026.04.07.717036

**Authors:** Areej Malik, Joshua R. Fletcher

## Abstract

Short-chain fatty acids (SCFAs) like butyrate and propionate are abundant microbiota-derived metabolites that influence bacterial physiology in host-associated niches like the gastrointestinal tract. However, their effects on *Staphylococcus aureus* under varying nutritional conditions remain incompletely understood. Here we investigated how SCFAs interact with glucose or galactose to regulate anaerobic growth, biofilm formation, and global transcription in *S. aureus*. Both SCFAs inhibit anaerobic growth in a dose-dependent manner but biofilm was differentially affected, with butyrate promoting and propionate suppressing biofilm formation. Glucose and galactose alleviated SCFA-mediated growth inhibition, with glucose exerting the strongest effect. Notably, glucose enhanced butyrate-associated growth and biofilm formation beyond glucose alone, whereas galactose produced more modest effects. Enzymatic and genetic analyses indicated that SCFA-sugar biofilms contain proteins and extracellular DNA and involve VraSR-dependent regulation. Transcriptomic profiling revealed broad metabolic reprogramming, including induction of urease genes, amino acid biosynthesis, and stress response pathways. Synergistic effects between butyrate and glucose were partially dependent on anaplerotic metabolism via pyruvate carboxylase, linking oxaloacetate regeneration from pyruvate to *S. aureus* SCFA adaptation. Together these findings demonstrate that the nutritional environment dictates whether SCFAs impair *S. aureus* growth or reprogram its physiology, promoting metabolic adaptation and biofilm formation under sugar-replete conditions.

## INTRODUCTION

*Staphylococcus aureus* is a Gram-positive opportunistic pathogen, and prominent biofilm-forming member of the phylum Firmicutes. Among the 52 recognized *Staphylococcus* species and 28 subspecies, *S. aureus* is the most clinically significant^1,2^. Approximately 20-30% of the human population is persistently colonized by *S. aureus*, most commonly in the nose and frequently in other body sites including skin, throat, axillae, groin, and intestinal tract^2,3^. Treatment of *S. aureus* infection has been increasingly complicated by the emergence and dissemination of antimicrobial resistance. Clinical isolates exhibit resistance to multiple classes of antimicrobial drugs, including ẞ-lactams, sulfonamides, tetracyclines, glycopeptides, and others^4^. Methicillin-resistant *Staphylococcus aureus* (MRSA) remains a leading example of drug-resistant bacteria and is a substantial public health burden, accounting for thousands of deaths annually in the United States^5–7^.

New interventions to selectively decolonize a host are needed to avoid further exacerbating antimicrobial resistance in *S. aureus* and other pathogens. One such approach is to increase the colonization resistance of the indigenous microbiota such that *S. aureus* is unable to colonize or persist, thereby decreasing the risk of severe infection. For example, probiotic-mediated clearance of *S. aureus* from the gut significantly diminishes its presence in the nasal sinuses^8^. This suggests that the gut may be a primary reservoir of *S. aureus* in humans, contributing to antibiotic associated diarrhea and heightened extraintestinal infection risk^9–11^. Indeed, nasal and intestinal colonization are correlated, suggesting a possible bidirectional transit mechanism^12^. Consistent with this model, intestinal *S. aureus* may also invade immune cells surveilling the luminal content of the gut and traverse intracellularly to extra-intestinal sites to initiate infections^8,13,14^. However, the mechanisms used by *S. aureus* to colonize the gut and metabolically integrate within the highly complex milieu of the gut microbiome are comparatively less well understood than for host communities of the skin or diseased airways.

In the gut, *S. aureus* likely encounters a highly competitive environment shaped by nutrient availability and microbiota-derived metabolites like short-chain fatty acids (SCFAs). These are metabolic byproducts of carbohydrate or amino acid fermentation, and are highly abundant in the gut^15,16^. While our group and others have shown that SCFAs impair *S. aureus* growth *in vitro*, it is still capable of colonizing and persisting in the gut, where SCFA concentrations are in the tens of mM^12,17–19^. The complex nutritional milieu of the gut likely exposes *S. aureus* to a variety of nutrients that could potentiate or mitigate the deleterious effects of SCFAs. For example, we have shown that the branched chain amino acid isoleucine promotes anteiso branched chain fatty acid production to partially restore growth of *S. aureus* in SCFA-rich media^19^. In a host, glycoprotein degradation by members of the gut microbiota can release sugars and peptides not only to the primary degrader but also nearby species, likely supporting their growth. Indeed, *S. aureus* can bind to intestinal mucus, placing it in close proximity to relevant glycan degraders^20^. Similarly, individuals with insulin resistance or diabetes, for whom *S. aureus* infection is of heightened concern, have elevated abundances of both *S. aureus* and sugars like glucose and galactose in their feces^21,22^. A clearer understanding of how the nutritional environment intersects with gut-relevant metabolites like SCFAs may reveal novel targets for therapeutic interventions that promote colonization resistance against *S. aureus*.

In this study, we examined the effects glucose or galactose on anaerobic growth and biofilm development by *S. aureus* in media with exogenous SCFAs. We show that both sugars alleviate SCFA-mediated anaerobic growth inhibition, albeit to different degrees, with butyrate synergizing with glucose to promote greater growth than glucose alone. We show that this is partially dependent upon the pyruvate carboxylase gene *pycA*, which converts pyruvate to oxaloacetate, as well as *ald1*, whose product mediates the reversible conversion of pyruvate to alanine. Together, these data reveal that access to glucose or galactose and their intermediate pyruvate within the gut may provide a metabolic advantage in the face of SCFA stress.

## MATERIALS AND METHODS

### Bacterial strains

*S. aureus* strain JE2 (plasmid-free derivative of USA300 LAC) was used as the wild type strain in all experiments. Mutants from the Nebraska Transposon Mutant Library were included where relevant^23^. The full list of strains and their specific transposon insertion identifiers can be found in **Supplemental File 1**.

### Growth curves

Cultures were prepared by inoculating the indicated *Staphylococcus aureus* strains into 2 mL of LB media and grown for 16-18 h at 37°C in a Coy anaerobic chamber (Coy). Cultures of NTML mutants included 4 *μ*g/mL erythromycin for selection. The overnight cultures were diluted 1:20 into LB media containing the indicated concentrations of sodium butyrate (Sigma Aldrich) sodium propionate (Sigma Aldrich), sodium pyruvate (Sigma Aldrich), urea (Sigma Aldrich), 0.5% w/v glucose (Sigma Aldrich), and 0.5% w/v galactose (Sigma Aldrich) in sterile tissue-culture treated 96-well plates (Genesee Scientific). A single concentration of 50 mM of each SCFA was used in all experiments with sugars. The plates were incubated without shaking at 37°C for 24h in a Cerillo microplate reader (Cerillo), which was set to record the optical density at 600 nm (OD_600_) every 30 min in the anaerobic chamber.

### Biofilm formation, visualization, and quantification

Overnight anaerobic cultures were diluted 1:20 in LB broth supplemented with the indicated concentrations of sodium butyrate, sodium propionate, sodium pyruvate, with and without 0.5% w/v glucose or galactose and incubated at 37°C in an anaerobic chamber. Biofilms were established in sterile, tissue-culture–treated 96-well plates with a total volume of 0.2 mL per well and incubated anaerobically without shaking at 37°C for either 24h or 48h. Following incubation, the plates were removed from the anaerobic chamber. The supernatant was carefully removed by pipetting, and the adhered biofilms were gently washed with 0.2 mL of 1X phosphate-buffered saline (PBS; Gibco). Biofilms were stained with 0.2 mL of 0.1 % crystal violet (Sigma Aldrich) for 10 min. After staining, the crystal violet was aspirated, and the stained biofilms were washed twice with 0.2 mL of sterile 1X PBS. The stained biofilms were then scraped and resuspended in 0.2 mL of 30% acetic acid (Fisher Scientific). Optical density at 570 nm was measured using a VarioSkan plate reader (Thermo Fisher Scientific) after shaking the plate at medium intensity for 15s.

### Proteinase K and DNase I treatment of 24h biofilms

As described above, static biofilms were established in sterile, tissue-culture–treated 96-well plates with a total volume of 0.2 mL per well and incubated anaerobically at 37°C for 24h. Following incubation and washes, the biofilms were treated with 0.2 mL of proteinase K (0.1mg/mL) in 20 mM Tris-HCl (pH 7.5) and 100 mM NaCl or 140 U/mL DNase I in 5 mM MgCl_2_. Plates were incubated statically at 37°C for 16h. After the treatments, the suspension was carefully removed by pipetting, the treated biofilms were then gently washed twice with 0.2 mL of 1X PBS, and stained with 0.2 mL of 0.1% crystal violet for 10 min. After staining, the crystal violet was aspirated, and the stained biofilms were washed twice with 0.2 mL of sterile 1X PBS. The stained biofilms were then scraped and resuspended in 0.2 mL of 30% acetic acid (Fisher Scientific). Optical density at 570 nm was measured using a VarioSkan plate reader after shaking the plate at medium intensity for 15s.

### RNA Extraction

Overnight anaerobic *S. aureus* JE2 cultures were diluted 1:20 into 3 mL of LB broth ± 50 mM sodium butyrate ± 0.5% w/v glucose in 15 mL conical tubes within an anaerobic chamber. Cultures were incubated for 24h at 37°C. After incubation, the cell suspensions were centrifuged at 4996 x g for 20 min at 4°C. The supernatant was aspirated without disturbing the cell pellet. Pellets were resuspended in 45 µL of LB media containing lysostaphin (Sigma Aldrich; final concentration 0.5 mg/mL) and incubated at 37°C for 20 min. TRIzol (1 mL; Invitrogen) was added, followed by incubation at room temperature for 5 min. Chloroform (0.2 mL; FisherScientific) was added and samples were vigorously mixed by hand for 15s, followed by incubation at room temperature for 5 min. Samples were then centrifuged at 12,000 rpm for 15 min at 4°C. The aqueous phase (∼0.5 mL) was transferred to RNase-free tubes containing equal volume of 95% EtOH (Koptec), mixed, and incubated for 5 min at room temperature. RNA was purified using the Zymo RNA Clean & Concentrator-5 kit (Zymo) with on-column DNase I treatment, according to the manufacturer’s instructions. Samples were stored at −80°C before they were sent for sequencing.

### RNA sequencing and analysis

Purified RNA was sequenced by SeqCenter (Pittsburgh, PA). Illumina’s Stranded Total RNA Prep Ligation with Ribo-Zero Plus kit was used to prepare libraries with unique 10 bp dual indices. Sequencing was performed on an Illumina NovaSeq X Plus to generate 150 bp paired end reads, which were demultiplexed via bcl-convert (v4.2.4), followed by adaptor trimming and quality control. Subsequently, the .fastq files were downloaded and transcripts were quasi-mapped and counted using Salmon^24^, with the coding sequences from the USA300 genome (https://aureowiki.med.uni-greifswald.de/Downloads) used to build the transcriptome index. The R package tximport (v1.30.0) was used to import Salmon output into RStudio for differential expression analysis using DESeq2 (v1.42.1)^25^. An adjusted p-value of <0.05 was considered statistically significant. Visualization of results was performed with the R packages ggplot2 (v4.0.1) and GraphPad Prism (v10.1.1).

### Statistical analyses

All data are presented as the mean ± standard error of the mean (SEM) for each experiment. Statistical analyses and data visualization were performed using GraphPad Prism 10 (v10.1.1). Quantitative experiments were conducted in technical duplicate and independently repeated at least three times.

### Data availability

All sequencing files are available at the NCBI Sequencing Read Archive (NCBI SRA) under the BioProject ID PRJNA1450098.

## RESULTS

### Dose-dependent inhibition of *S. aureus* anaerobic growth by short-chain fatty acids

To evaluate the impact of short-chain fatty acids (SCFAs) on *S. aureus* anaerobic growth, the MRSA strain JE2 was cultured in LB medium supplemented with increasing concentrations of (0-100mM) of sodium butyrate (NaB) or sodium propionate (NaP) and OD_600_ was monitored over 24h every 30 minutes. The untreated control exhibited robust growth, characterized by a rapid exponential phase and entry into the stationary phase by approximately 6 hours. In contrast, exposure to 10 mM and 25 mM of either SCFA resulted in reduced growth rates and lower maximum OD_600_ values relative to the untreated control (**Fig. 1A, 1D**). At higher concentrations, 50 mM and 100 mM, growth was markedly suppressed throughout the 24h incubation period. Quantitative analysis of the area under the curve (AUC) supported these findings, showing a significant concentration-dependent decline (**Fig. 1B, 1E**). These results demonstrate that butyrate and propionate inhibit *S. aureus* proliferation in a dose-dependent manner under anaerobic conditions, consistent with what has been reported for aerobic growth^17^.

**Figure 1.**
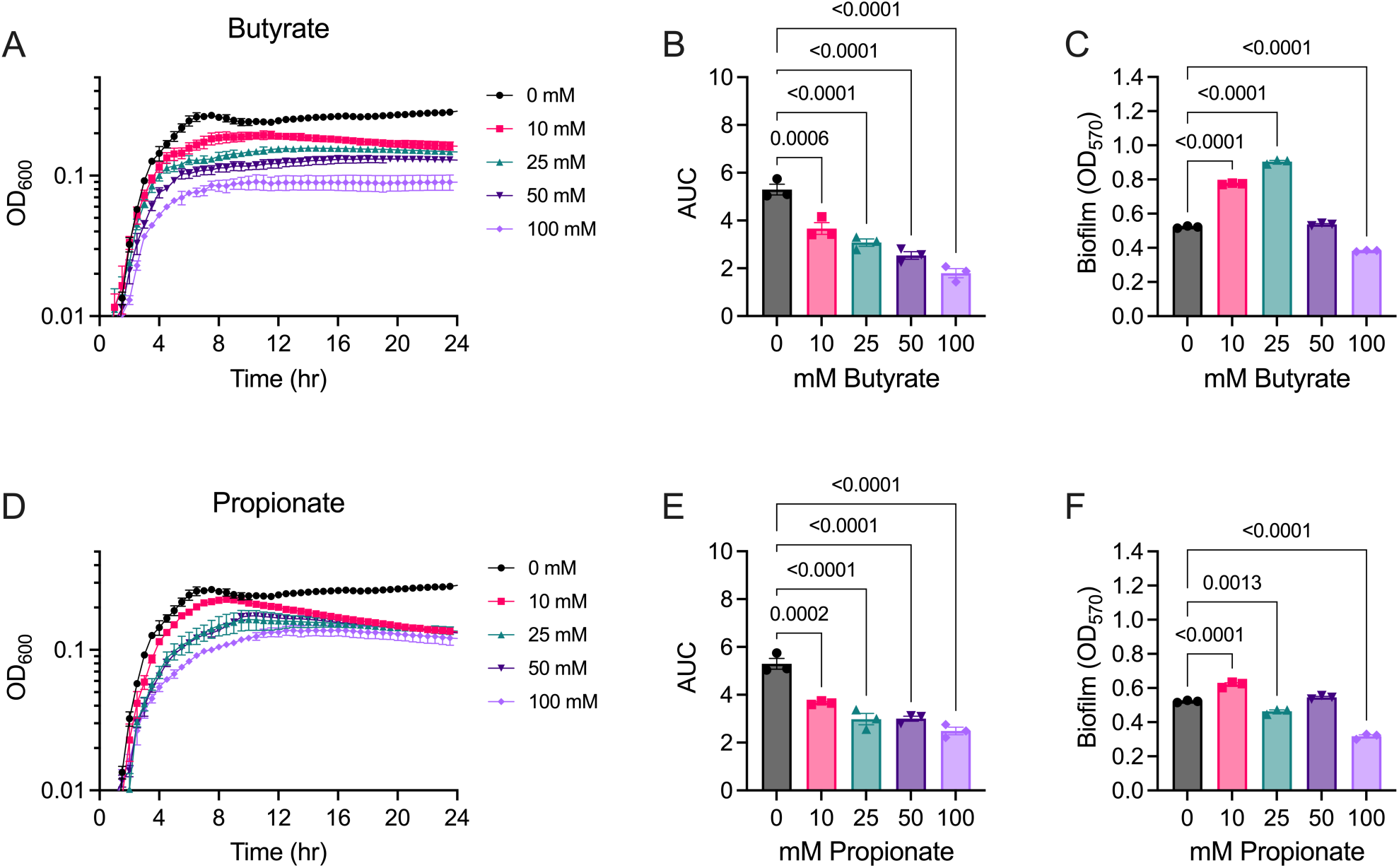
SCFAs inhibit *Staphylococcus aureus* JE2 anaerobic growth in a concentration-dependent manner. **A and D)** Anaerobic growth curves of *S. aureus* JE2 in LB medium supplemented with the indicated concentrations of butyrate or propionate. **B and E)** Area under the curve analyses of the growth curves in A and D. **C and F)** Biofilm formation after 24h incubation in LB supplemented with the indicated concentrations of butyrate or propionate, as measured by crystal violet staining. Data represent the mean ± standard error of the mean of three independent biological replicates. P-values in B, C, E, and F derive from an ordinary one-way ANOVA with Dunnett’s correction for multiple comparisons, with a single pooled variance. All conditions were compared to the 0 mM condition.

### Opposing effects of short-chain fatty acids on *S. aureus* biofilm formation

Treatment with SCFAs also altered anaerobic biofilm formation by JE2 in a dose-dependent manner (**Figs. 1C and 1F**). Growth with butyrate lead to a marked enhancement of 24h anaerobic biofilms at 10 and 25 mM relative to the untreated control, while 50 and 100 mM were comparable to the untreated biofilms. The most pronounced effect was observed at ≥25 mM after 48h of incubation (**Suppl. Fig. 1A**). Biofilm accumulation in media supplemented with butyrate was consistently higher at 48h compared to 24h across all concentrations, showing sustained development over time. In contrast, propionate mostly inhibited biofilm formation, although there was a modest increase at 10 mM at the 24h time point relative to untreated controls. A significant reduction in OD_570_ was observed at all propionate concentrations at or above 25 mM. The decrease in biofilm mass was evident at both 24 and 48h, indicating that propionate exerts a largely suppressive effect (**Suppl. Fig. 1B**). Together, these findings show that SCFAs differentially modulate *S. aureus* biofilm formation in a time- and concentration-dependent manner.

### Glucose and galactose relieve *S. aureus* growth impairment by SCFAs

As a facultative anaerobe, *S. aureus* can fuel rapid growth via glycolysis in oxygen-limited environments, therefore we examined if glucose supplementation allowed it to overcome SCFA-mediated growth inhibition. Similarly, the environments where *S. aureus* would encounter SCFAs (CF airways or mammalian gut) may also have elevated levels of monosaccharides like galactose due to the activity of glycan foragers in the microbiota, e.g., the *Streptococcus*, *Segatella*, and *Bacteroides* genera^26–29^, therefore we included galactose in our growth assays. As expected, both sugars supported robust anaerobic growth of JE2 relative to LB alone and markedly enhanced growth in the presence of both SCFAs (**Fig. 2A, B, D, E**). Surprisingly, the combination of butyrate and glucose resulted in greater growth than glucose alone, evidenced by higher OD_600_ after 24h of growth and higher AUC. While galactose partially enhanced growth relative to cultures with each SCFA alone, the effect was not as pronounced as with glucose. Indeed, the AUCs of SCFA cultures supplemented with galactose were lower than galactose-only cultures. Collectively, these data demonstrate that while both sugars enhance *S. aureus* growth in the presence of SCFAs, glucose was more effective.

**Figure 2.**
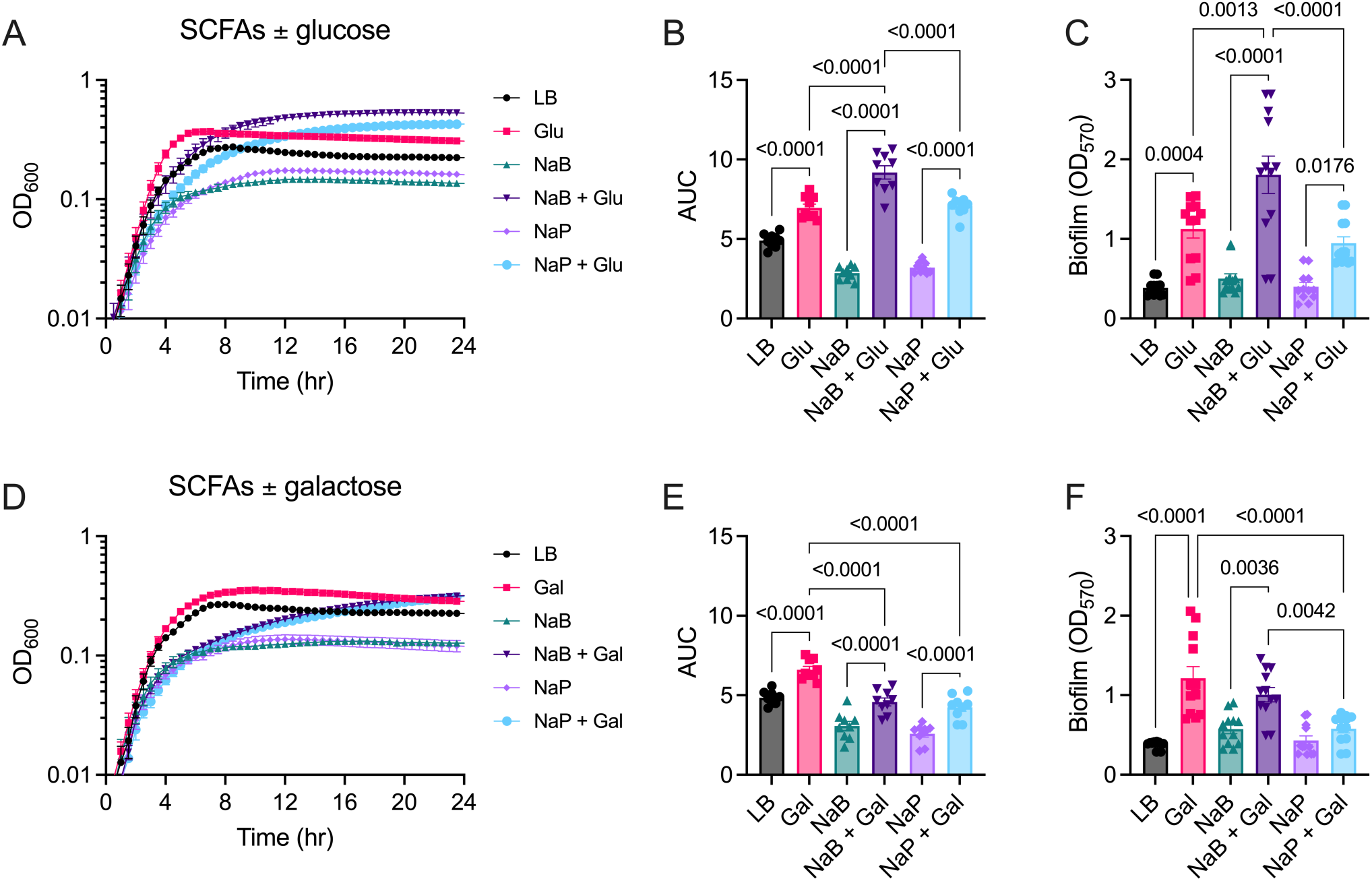
SCFA-mediated inhibition of *Staphylococcus aureus* JE2 anaerobic growth is relieved by sugar supplementation. **A and D)** Growth curves of *S. aureus* JE2 in LB ± 50mM SCFAs ± 0.5% w/v glucose or galactose. **B and E)** Area under the curve analysis of the growth curves in A and D. **C and F)** Biofilm formation after 24h incubation in LB ± 50mM SCFAs ± 0.5% w/v glucose or galactose, as measured by crystal violet staining. Data represent the mean ± standard error of the mean of at least nine independent biological replicates. P-values are from an ordinary one-way ANOVA with Tukey’s correction for multiple comparisons, with a single pooled variance. All conditions were compared to each other, though only relevant comparisons are shown for clarity.

### Biofilm formation by *S. aureus* is promoted by the combination of butyrate and glucose

SCFAs differentially modulated biofilm formation in a sugar- and time-dependent manner (**Fig. 2C, 2F, Suppl. Fig. 2**). Supplementation of LB medium with either glucose or galactose resulted in comparable biofilm biomass at 24h, as expected. In SCFA media, the addition of glucose enhanced biofilm biomass relative to LB and SCFA-only controls at 24h and 48h. While the data were variable, the combination of butyrate and glucose produced the highest levels of biofilm accumulation at both time points, exceeding glucose-only controls and propionate plus glucose. This indicates a stronger biofilm-promoting effect of butyrate with glucose that is consistent with enhanced growth in this condition. Conversely, galactose supplementation of the SCFA media resulted in intermediate biofilm formation that was greater than LB or SCFA-only controls, but lower than the galactose-only condition. For both butyrate- and propionate-treated cultures, biofilm biomass was consistently higher in glucose-supplemented media than in galactose-supplemented media at both 24h and 48h. Together, these data indicate that biofilm development in *S. aureus* is strongly influenced by the interaction between sugar type and SCFAs, with glucose preferentially supporting SCFA-driven biofilm accumulation and butyrate acting as an enhancer.

### Cell wall stress signaling contributes to butyrate + sugar-mediated anaerobic biofilms

Given the effect of each SCFA on sugar-mediated biofilm formation, we sought to identify potential mechanisms driving these phenotypes to determine if biofilm composition varied across conditions. Staphylococcal biofilm matrices are composed of extracellular polysaccharides, proteins, and extracellular DNA, although polysaccharides are reported to be less prominent in MRSA isolate biofilms^30^. We screened six transposon mutants with genetic links to biofilms to identify plausible mechanisms by which the combination of SCFAs and sugars contribute to surface attachment or biofilm structure (**Fig. 3**). These mutants were chosen for loss of extracellular polysaccharide (*icaA*::tn)^31^, loss of cell wall protein anchoring (*srtA*::tn)^32^, protease overproduction (*sarA*::tn)^33^, alteration of the stringent response (*relP*::tn)^34^, and loss of cell wall stress sensing and response (*vraR*::tn, *vraS*::tn). The *vraSR* mutants were included for two reasons: i) they are linked to intermediate vancomycin resistance in *S. aureus* and in a previous study, we demonstrated that SCFAs modestly protected *S. aureus* from vancomycin *in vitro*^19^, and ii) a double *vraSR* mutant is reported to form less biofilm^35^.

**Figure 3.**
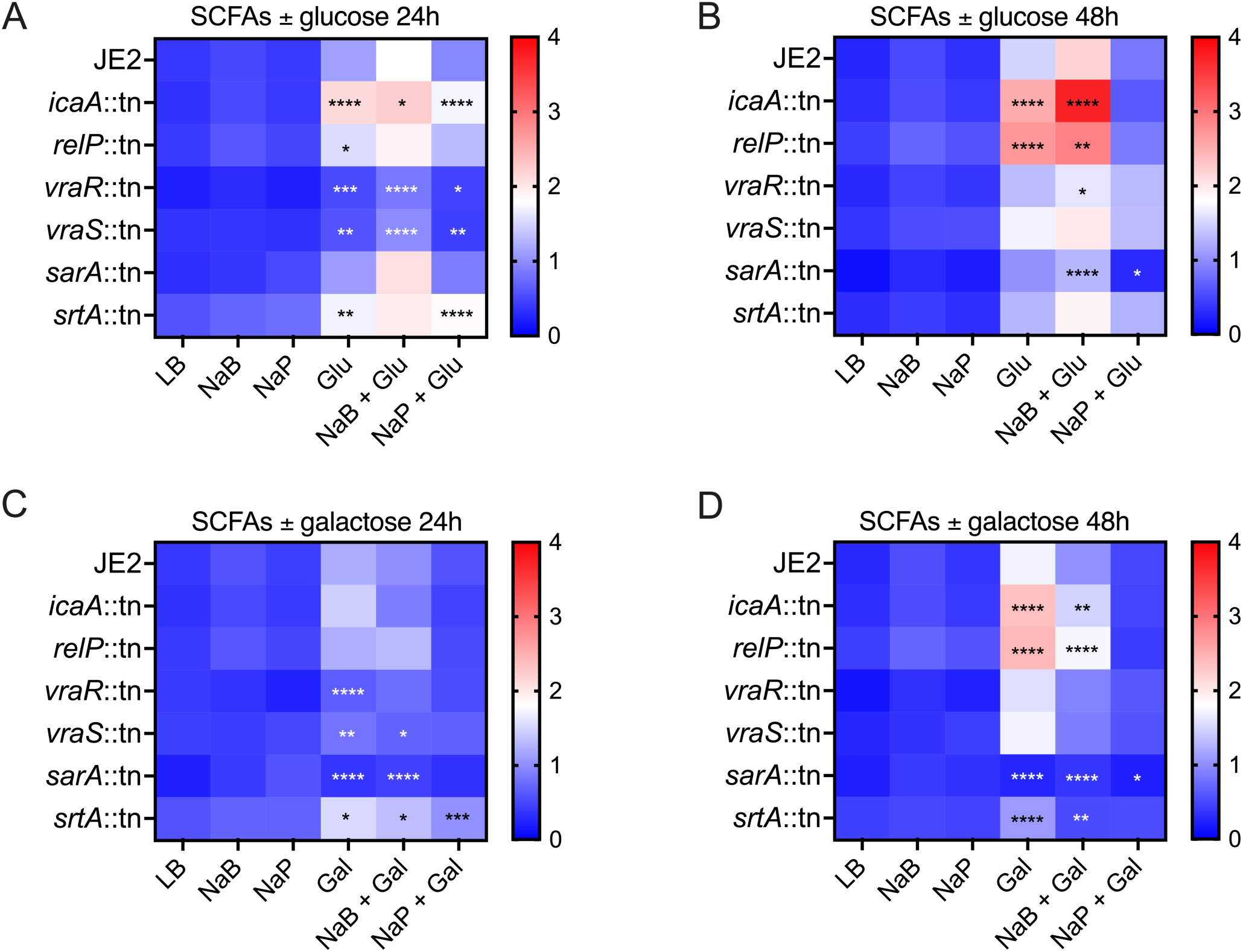
Combined SCFA and sugar supplementation drives time- and mutant-dependent *Staphylococcus aureus* biofilm responses. Heatmaps of biofilm OD_570_ from wild type *S. aureus* JE2 and transposon mutants cultured in LB medium supplemented with 50 mM SCFAs in the absence or presence of 0.5% w/v glucose after **A)** 24h or **B)** 48h incubation, or 0.5% w/v galactose after **C)** 24h or **D)** 48h incubation. Columns denote media conditions and rows denote strains. Blue indicates low biofilm formation and red indicates enhanced biofilm formation. Data represent the mean ± standard error of the mean of at least three independent biological replicates. P-values are from an ordinary two-way ANOVA with Dunnett’s correction for multiple comparisons, with a single pooled variance. NS (no asterisk) P > 0.05, * P < 0.05, ** P < 0.01, *** P < 0.001, **** P < 0.0001. All mutant strains were compared to wild type JE2 within a given medium.

We repeated 24 and 48h biofilm assays in LB supplemented with SCFAs and glucose or galactose. The *icaA*::tn and *relP*::tn mutants had elevated biofilms in SCFA media with either sugar at both 24 and 48h, although the mechanisms underlying these phenotypes are unclear. Surprisingly, loss of *srtA* lead to increased biofilm formation in both sugar ± SCFA media at 24h, though biofilm biomass at 48h was comparable to wild type JE2. Both *vraS*::tn and *vraR*::tn formed less biofilm at 24h in these media, consistent with our previously published data implicating cell wall stress in response to SCFAs during aerobic growth^19^. The 24h biofilms from the *sarA*::tn mutant in LB supplemented with glucose and SCFAs were similar to those from JE2, but significantly lower in media containing galactose. By 48h, the *sarA*::tn biofilms were lower in media with either sugar plus SCFAs, consistent with the overproduction of proteases degrading the proteinaceous elements of the biofilm architecture. We also characterized their growth in each medium and found that *relP*::tn had a small but significant reduction in AUC in butyrate plus glucose medium relative to JE2, while *sarA*::tn, *vraR*::tn, and *vraS*::tn grew less well than JE2 in propionate plus glucose **(Suppl Fig. 3)**. Conversely, *srtA*::tn grew better than JE2 in butyrate-only medium, while *sarA*::tn grew worse than JE2 in galactose medium and its growth was further repressed in galactose plus SCFA media.

### Butyrate plus glucose biofilms are more susceptible to DNase I than glucose-only biofilms

Given the enhanced growth and biofilm formation of JE2 in LB with butyrate and glucose, we narrowed our focus to this condition and complemented our genetic approach with proteinase K or DNase I treatments to obtain a course-grained approximation of matrix composition (**Suppl. Fig 4**). In glucose-supplemented medium, proteinase K significantly reduced biofilm biomass, indicating a strong dependence on proteinaceous matrix components, whereas DNase I did not have an effect. Conversely, both Proteinase K and DNase I degraded the butyrate plus glucose biofilms, indicating that the abundance of extracellular DNA may increase when butyrate and glucose are present. Taken together with biofilm mutant data from Figure 3, these results indicate that biofilms formed in the presence of butyrate and glucose rely on deposition of both protein and extracellular DNA.

### Butyrate plus glucose significantly alters the *S. aureus* anaerobic transcriptome

The enhanced growth of *S. aureus* in media with butyrate and glucose suggests a metabolic reprogramming that minimizes the deleterious effects of butyrate. We performed RNA seq to identify genes and pathways that may be implicated in this phenotype (**Fig. 4, Suppl. Fig 5, Suppl. File 1**). We grew *S. aureus* JE2 in LB with and without butyrate, glucose, or both for 24h in anaerobic conditions and compared their transcriptional profiles. Principal component analysis (PCA) demonstrated clustering by growth condition, indicating that media supplementation is a major driver of global transcriptional variation. Importantly, the butyrate plus glucose samples formed a discrete cluster rather than overlapping with either glucose or butyrate alone, suggesting a combinatorial effect on the *S. aureus* transcriptome driven by each **(Fig. 4A)**. Consistent with this, several hundred genes were significantly differentially expressed at an adjusted p-value of less than 0.05 across every comparison.

**Figure 4.**
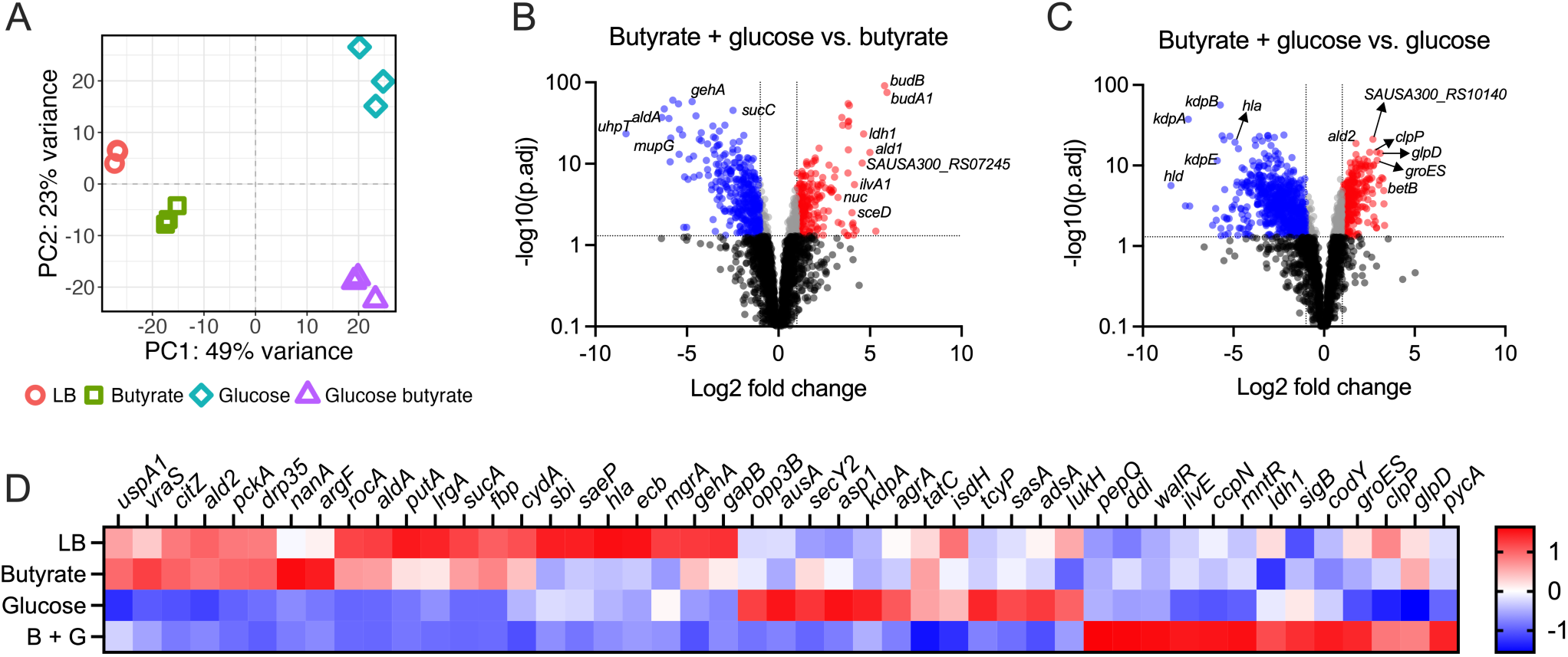
Global transcriptional responses of *S. aureus* to butyrate and glucose. **A)** Principal component analysis (PCA) of *S. aureus* JE2 transcriptomic profiles showing clustering of samples grown in LB only (salmon circles), butyrate only (green squares), glucose only (teal diamonds), and glucose plus butyrate (purple triangles). **B)** Volcano plot comparing JE2 gene expression in butyrate plus glucose versus butyrate alone. **C)** Volcano plot comparing JE2 gene expression in butyrate plus glucose versus glucose alone. Differentially expressed genes are indicated by a minimum of two-fold change in expression at an adjusted p-value < 0.05, with significantly upregulated transcripts colored red and significantly downregulated transcripts colored blue. **D)** Heatmap of z-scores of normalized read counts for selected *S. aureus* genes across the indicated media. The color scale represents relative expression (red, higher; blue, lower).

Exposure to butyrate alone resulted in extensive transcriptional remodeling compared to LB, with genes associated with urease activity (*ureC, ureB,* and *ureA*)^36^ and stress response pathways (*ctsR, groES, clpP*)^37^ among the most significantly upregulated, while amino acid biosynthesis genes (*argG, argH, thrS*) were downregulated^38,39^. Additionally, increased expression of genes such as *mccA* and *mccB* points to a shift toward cysteine metabolism^40^. Conversely, several virulence-associated genes (*hla* and *lukH*) were downregulated, along with a broader set of genes involved in metabolic versatility and nutrient acquisition^41,42^. These include genes linked to carbohydrate and fermentative metabolism (*ldh1*) and nucleotide and cofactor biosynthesis (*gua*), suggesting a contraction of biosynthetic capacity under butyrate conditions^43,44^. Downregulation of these pathways indicates reduced reliance on diverse nutrient sources and a shift away from energy-intensive biosynthetic processes. Together, these data suggest that butyrate drives a distinct physiological state characterized by stress responses and nitrogen metabolism, coupled with suppression of virulence and a narrowing of metabolic flexibility **(Fig. 4B, D)**.

The addition of glucose to butyrate led to a broader and more pronounced transcriptional response compared to butyrate alone. A large number of genes exhibited significant differential expression with higher magnitude fold changes, indicating that glucose strongly modulates butyrate-affected pathways. Notably, genes involved in carbohydrate metabolism and fermentation (e.g., *ldh1* and *ald1*) were significantly upregulated, consistent with the expected metabolic shift towards glycolysis and fermentative processes^44,45^. Urease-associated genes (*ureC, ureD, ureE,* and *ureF*) again showed strong upregulation, suggesting enhanced nitrogen metabolism under combined conditions. In contrast, genes involved in alternative nutrient utilization and metabolic flexibility were downregulated, including those associated with amino acid metabolism (*rocF, putA*)^46,46^, sialic acid utilization (*nanA, nanT*)^47^, TCA cycle/energy metabolism (*sucC*)^48^, and lipid metabolism (*gehA*)^49^, along with additional metabolic enzymes such as *aldA*^50^. Compared to JE2 grown in glucose, the glucose plus butyrate samples exhibited robust induction of stress response genes (*groES, clpP*) while downregulating virulence factors like *hla* and *hld*^51^, as well as the quorum-regulated *kdpDE* two component system and associated potassium transport system encoded by *kdpFABC*^52^. These findings indicate that glucose not only overrides aspects of the butyrate response but also modifies fermentative, stress, and ionic homeostasis pathways **(Fig. 4C, D)**.

Collectively, these data demonstrate that butyrate and glucose exert independent and combinatorial effects on the *S. aureus* JE2 transcriptome. Butyrate induces a transcriptional program consistent with metabolic adaptation and attenuation of expression of specific virulence genes, whereas glucose promotes fermentative metabolism and potentiates key adaptive pathways, including nitrogen metabolism and stress response pathways. The distinct transcriptional landscape observed under combined conditions reveals the importance of metabolite interactions in shaping bacterial physiology and highlights how nutrient availability may influence pathogenic potential in complex host-associated environments like the gut.

### Pyruvate-driven oxaloacetate and alanine synthesis relieves butyrate inhibition of *S. aureus* growth

Our transcriptomics revealed a distinct set of genes that were uniquely sensitive to the presence of both butyrate and glucose. To link gene expression to the growth enhancement we observed in butyrate plus glucose medium, we screened several transposon mutants in genes that displayed elevated expression (**Fig. 5**). As expected, wild-type JE2 exhibited robust growth in butyrate plus glucose medium, while two mutants displayed impaired growth (*pycA*::tn and *ald1*::tn), indicating a critical role for these pathways in supporting growth under these conditions. PycA converts pyruvate to oxaloacetate, while Ald1 catalyzes the reversible conversion of pyruvate to alanine, suggesting that pyruvate metabolism is central to the robust growth of *S. aureus* in conditions where butyrate and glucose are present. In contrast, *budB*::tn, *norB*::tn, *ureD*::tn, and *clpC*::tn mutants exhibited no defects in growth compared to JE2, despite robust induction of their transcripts in this medium. Although, the *ureD*::tn mutant did not have a phenotype, urea is not a component of LB, therefore it would not be in abundance for the urease proteins to act upon. Urease activity has been shown to promote acid resistance in a skin-like medium by raising pH^38^, therefore we assessed whether urea supplementation would rescue growth of wild type JE2 when butyrate was present, but it failed to do so (**Suppl. Fig. 6**). The requirement for *ald1* and *pycA* genes for optimal growth suggests that this transcriptional reprogramming may be an adaptive strategy, allowing *S. aureus* to thrive in nutrient-rich environments where organic acids and fermentable sugars co-exist.

**Figure 5.**
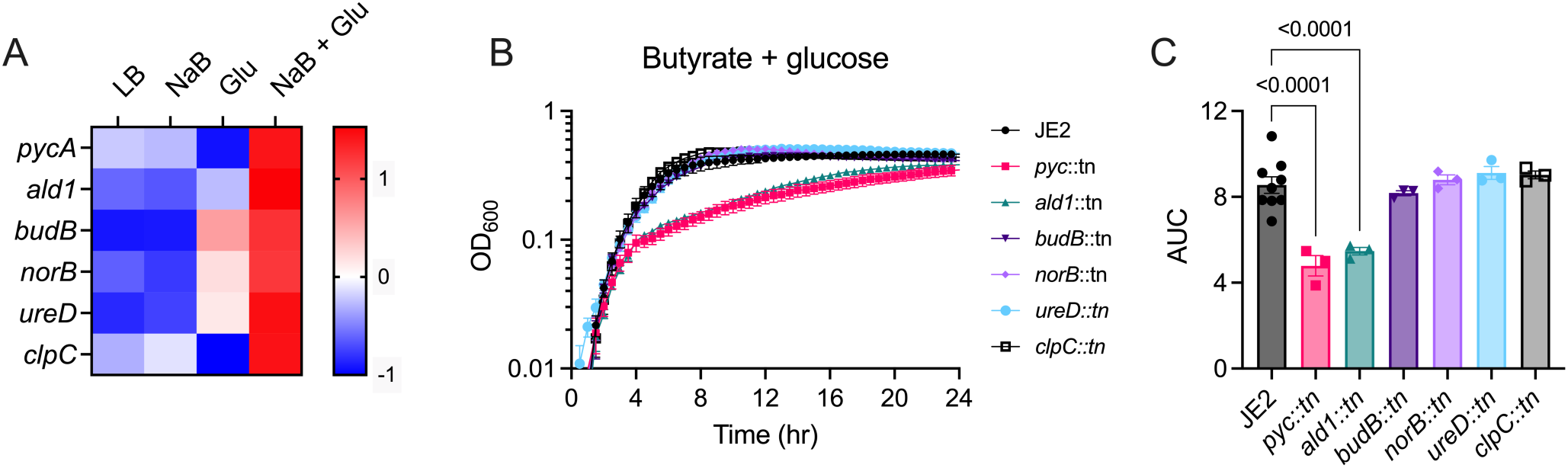
Pyruvate metabolism drives enhanced growth of *S. aureus* in butyrate- and glucose-containing medium. **A)** Z-scores of normalized read counts of the indicated genes across all four media conditions used in RNA seq analyses. **B)** Growth curves of JE2 and transposon mutants of the genes from A in LB with 50 mM butyrate and 0.5% (w/v) glucose. **C)** Area under the curves (AUC) from the growth curves in B. Data represent the mean and error bars are the standard error of the mean of at least three independent biological replicates. P-values are from an ordinary one-way ANOVA with Dunnett’s correction for multiple comparisons, with a single pooled variance. All mutants were compared to JE2.

To support the hypothesis that pyruvate was responsible for alleviation of butyrate-mediated growth inhibition, JE2 was cultured in butyrate medium with various concentrations of pyruvate (**Fig. 6**). As expected, butyrate alone significantly limited bacterial growth. Supplementation with pyruvate led to robust restoration of growth, with 2.0% pyruvate producing the most substantial effect. This rescue was evident as higher optical densities compared to butyrate alone and in area under the curve analysis. We then screened the *ald1*::tn and *pycA*::tn mutants in butyrate medium with 2% pyruvate (**Fig. 7**). The *ald1*::tn mutant was more sensitive to growth inhibition by butyrate than JE2 but grew similar to the wild type in LB supplemented with pyruvate. It displayed a modest growth defect in pyruvate-supplemented butyrate medium. As the Ald1 protein can convert pyruvate to alanine and vice versa, we interpret this to mean that the former reaction is needed for full growth when butyrate is present, as the excess pyruvate in this medium would mask the mutant’s deficit for the latter reaction. The *pycA*::tn mutant was also more sensitive to butyrate alone than JE2, although it did not reach statistical significance (p = 0.0565). Though pyruvate alone also boosted the growth of *pycA*::tn, its AUC was modestly reduced compared to JE2. When grown in butyrate medium supplemented with pyruvate, this mutant was significantly less fit, reaching a lower terminal OD600 with less than half the AUC of JE2. Together, these findings indicate that efficient anaerobic growth of *S. aureus* in the presence of butyrate and glucose depends on anaplerotic regeneration of oxaloacetate and biosynthesis of alanine.

**Figure 6.**
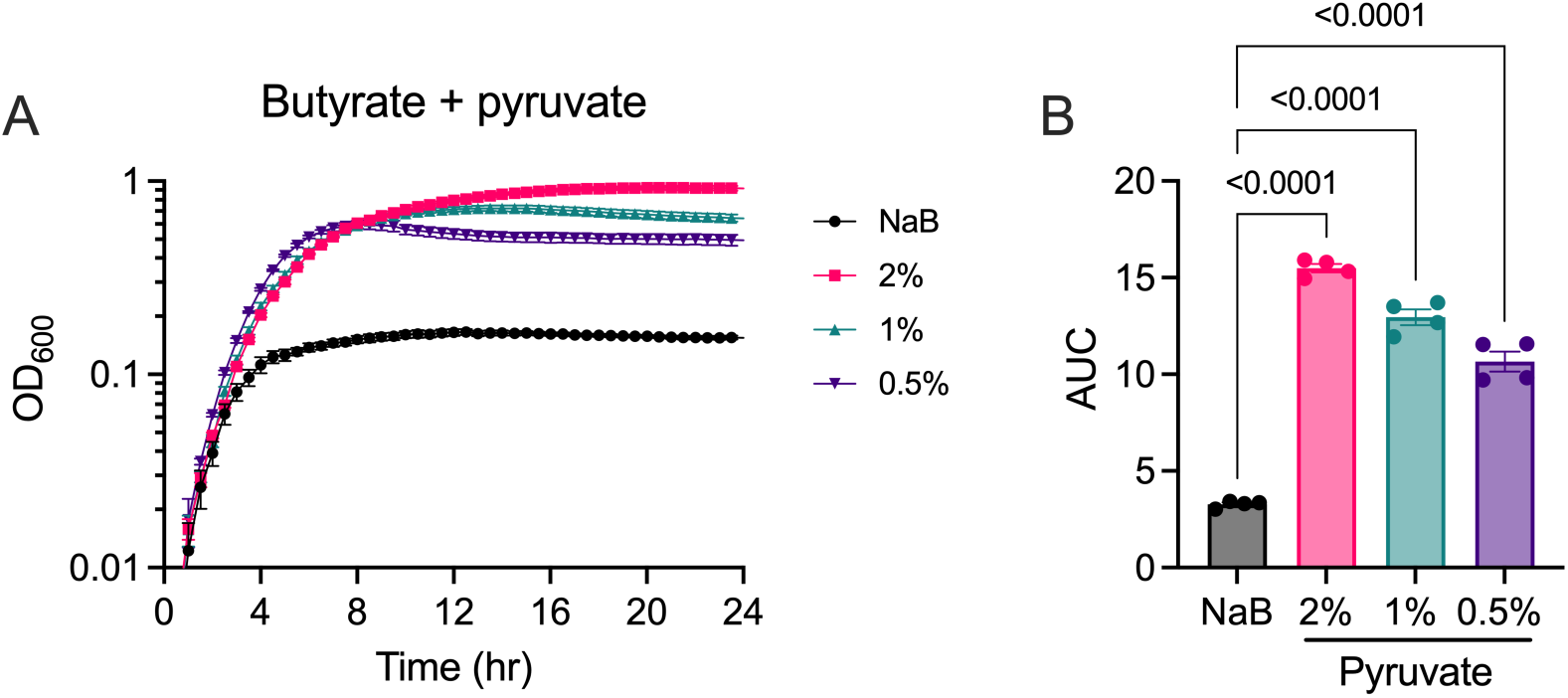
Pyruvate supplementation allows *S. aureus* to overcome butyrate-mediated growth inhibition. **A)** Growth curves of JE2 in LB medium with 50 mM butyrate and the indicated concentrations (w/v) of pyruvate. **B)** Area under the curve analyses of the growth curves in A. Data represent the mean and error bars are the standard error of the mean of four independent biological replicates. P-values are from an ordinary one-way ANOVA with Dunnett’s correction for multiple comparisons, with a single pooled variance. All conditions were compared to NaB.

**Figure 7.**
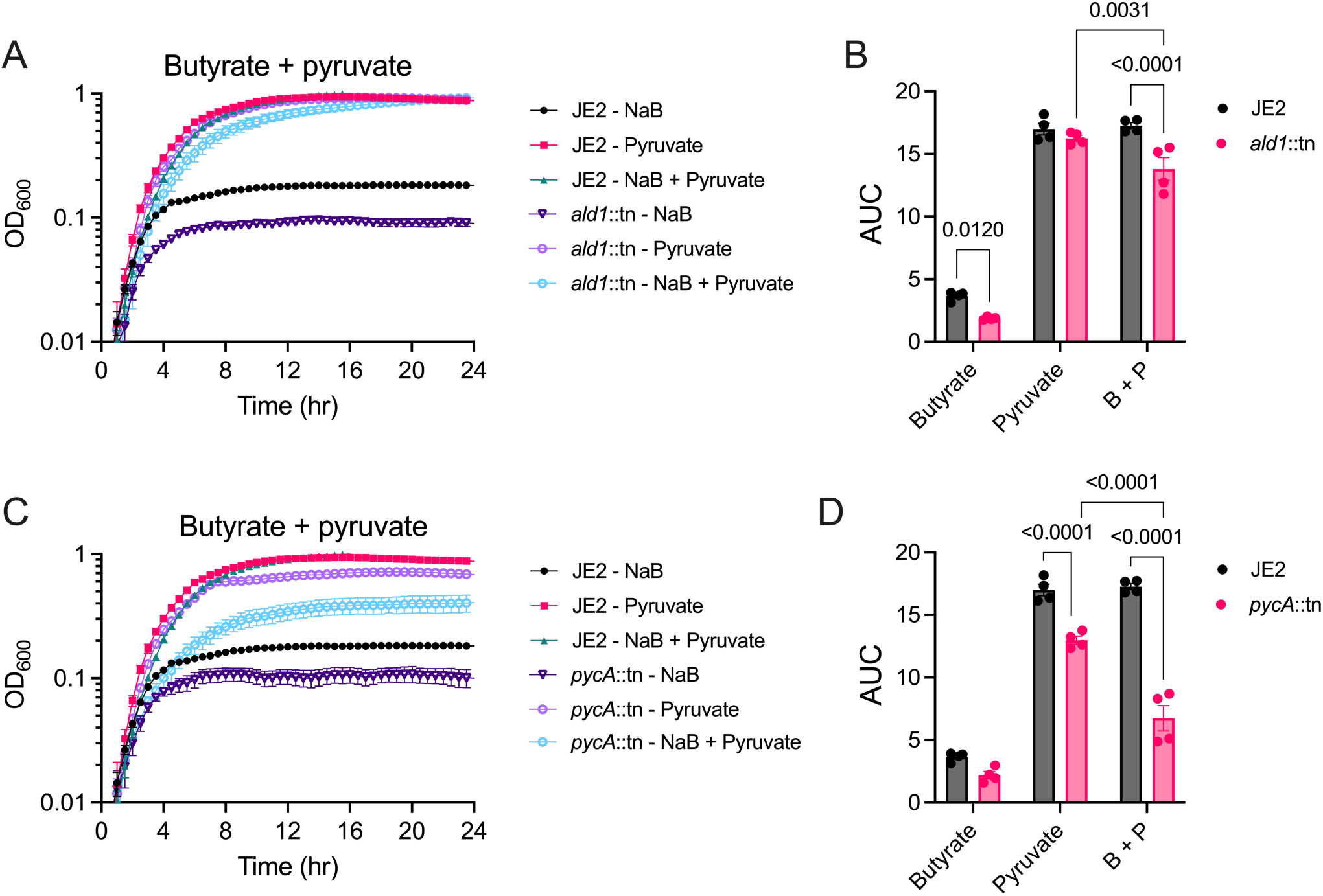
Alanine synthesis and anaplerotic regeneration of oxaloacetate are needed for pyruvate to alleviate butyrate-mediated growth inhibition of *S. aureus*. **A and C)** Growth curves and **B and D)** AUC analyses of JE2 and the *ald1*::tn and *pycA*::tn mutants in LB supplemented with 50 mM butyrate with and without 2% (w/v) pyruvate. Data represent the mean and error bars are the standard error of the mean of four independent biological replicates. P-values in B and D are from an ordinary two-way ANOVA with Tukey’s multiple comparisons test, with a single pooled variance. For clarity, only the relevant comparisons are shown.

## DISCUSSION

In this study, we demonstrate that short-chain fatty acids (SCFAs) exert complex, context-dependent effects on *Staphylococcus aureus* physiology under anaerobic conditions, simultaneously constraining planktonic growth while reshaping biofilm formation, metabolism, and gene expression. Our findings reveal that these effects are not only concentration-dependent but are profoundly influenced by available sugars, highlighting the importance of metabolic context in determining bacterial behavior that could be relevant in host-associated environments. A key finding of this study is that exogenous sugars, particularly glucose, can override SCFA-mediated growth inhibition and enhance biofilm formation. Glucose supplementation not only restored growth in the presence of SCFAs but, in combination with butyrate, resulted in even greater growth. This unexpected effect suggests that butyrate induces a response that synergizes with the effects of high glycolytic flux, mitigating its inhibitory effects. Indeed, several stress response pathways were induced in butyrate plus glucose medium relative to either supplement alone. Galactose, while also supporting growth, was less effective than glucose, indicating that the efficiency or regulation of carbon utilization pathways plays a critical role in determining the outcome of SCFA exposure. Biofilms from galactose plus SCFA media were attenuated compared to glucose conditions, suggesting that not all carbon sources equally support extracellular matrix production or maturation. These findings are particularly relevant to host environments such as the gut, where both SCFAs and sugars are present and likely fluctuate based on diet and microbiota activity^22^. Under such conditions, *S. aureus* may exploit available nutrients to overcome inhibitory metabolites, persisting through enhanced growth and biofilm formation.

Despite suppressing planktonic growth, SCFAs exerted divergent effects on biofilm formation. Butyrate enhanced biofilm formation in a time- and concentration-dependent manner, whereas propionate was largely inhibitory. This dichotomy suggests that each SCFA engages different regulatory or metabolic pathways that influence biofilm formation. The ability of butyrate to promote biofilm formation despite limiting growth is particularly notable and implies a shift toward a sessile, potentially more resilient lifestyle under metabolic stress. In contrast, the inhibitory effect of propionate on biofilm formation may reflect a stronger disruption of pathways required for matrix production or cellular adhesion to surfaces and other cells. Genetic analysis supported a role for protein-mediated biofilm structure, particularly implicating the VraSR two-component system. The reduced biofilm formation observed in *vraR::tn* and *vraS::tn* mutants at early time points, combined with the sensitivity of biofilms to protease treatment, suggests that VraSR-regulated factors like adhesins or other surface-associated proteins are critical for biofilm establishment under butyrate and glucose conditions. The phenotype of the *sarA::tn* mutant at later time points is consistent with protease-mediated degradation of matrix proteins, reinforcing the importance of protein stability in maintaining biofilm integrity. Interestingly, the increased biofilm formation observed in the *icaA::tn* mutant suggests that polysaccharide-independent mechanisms can compensate under certain metabolic conditions, highlighting the plasticity of *S. aureus* biofilm composition.

RNA-seq analysis revealed extensive metabolic reprogramming in response to SCFAs and glucose. Butyrate alone induced a transcriptional profile consistent with stress adaptation, including upregulation of urease genes that may contribute to pH homeostasis and nitrogen metabolism. Concurrent downregulation of virulence factors suggests that butyrate promotes a physiological state prioritizing survival over acute pathogenicity. The distinct clustering in the PCA analysis indicates that the response to butyrate and glucose likely represents a unique physiological state. Similarly, upregulation of the *vra* operon supports a role for cell envelope stress responses in adapting to SCFA-rich environments. These transcriptional changes likely underpin the enhanced growth and biofilm phenotype that we observed and provide insight into the mechanisms by which *S. aureus* integrates environmental signals to optimize survival.

Building on our transcriptomic analysis, validation of mutants in key butyrate- and glucose-responsive genes further supports the model for metabolic reprogramming underlying the enhanced growth phenotype. The induction of *ald1* and *pycA* transcripts under the combined butyrate and glucose condition, coupled with the pronounced growth defects observed in the corresponding transposon mutants, underscores the importance of pyruvate flux and anaplerotic metabolism in this context. The impaired growth of the *pycA::tn* mutant in particular highlights the necessity of oxaloacetate regeneration, although TCA cycle genes were downregulated under these conditions. This suggests that oxaloacetate is needed for other physiological processes. Similarly, the mildly reduced growth phenotype of the *ald1::tn* mutant indicates a role for alanine dehydrogenase in converting pyruvate to alanine, which may be needed to counteract impairment of peptidoglycan crosslinking, as has been shown due to acetate stress^53^. In contrast, the lack of growth defect in the *ureD::tn* mutant suggests that urease activity is not essential for growth in this medium and instead reflects a generalized response to acid stress imposed by butyrate. Consistent with this phenotype, supplementation of butyrate medium with urea did not rescue growth.

Together, our findings support a model in which SCFAs act not only as growth-inhibitory metabolites, but also as signaling cues that drive extensive metabolic and physiological reprogramming in *S. aureus*, with outcomes that are highly dependent on nutrient context. In anaerobic environments enriched in fermentable sugars, *S. aureus* can overcome SCFA-mediated stress by redirecting carbon flux through key metabolic nodes, including pyruvate metabolic pathways, to support growth and promote biofilm formation. The requirements for enzymes such as Ald1 and PycA highlights the importance of metabolic flexibility in enabling this adaptation, while the induction of stress-associated pathways further underscores the coordinated response to combined acid stress and nutrient signals. These findings suggest that *S. aureus* may leverage SCFAs in its environment as cues to transition towards a metabolically optimized, biofilm-associated state that could enhance persistence in polymicrobial communities commonly found in hypoxic or anaerobic environments like the cystic fibrosis or chronic rhinosinusitis airways, or the mammalian gut. Indeed, in this and previous studies, butyrate induced expression of the *nan* genes, whose products import and metabolize sialic acid^19,47^. *S. aureus* lacks a sialidase to liberate sialic acids, but many anaerobic members of mucus-rich diseased airways or gut communities do encode sialidases, and thus their SCFA byproducts may prime *S. aureus* to benefit from this enzymatic activity. More broadly, this work has important implications for understanding *S. aureus* behavior in polymicrobial and host-associated niches, where fluctuating metabolite landscapes and nutrient availability shapes pathogen physiology and may drive colonization and infection.

---

**Supplemental Figure 1.**
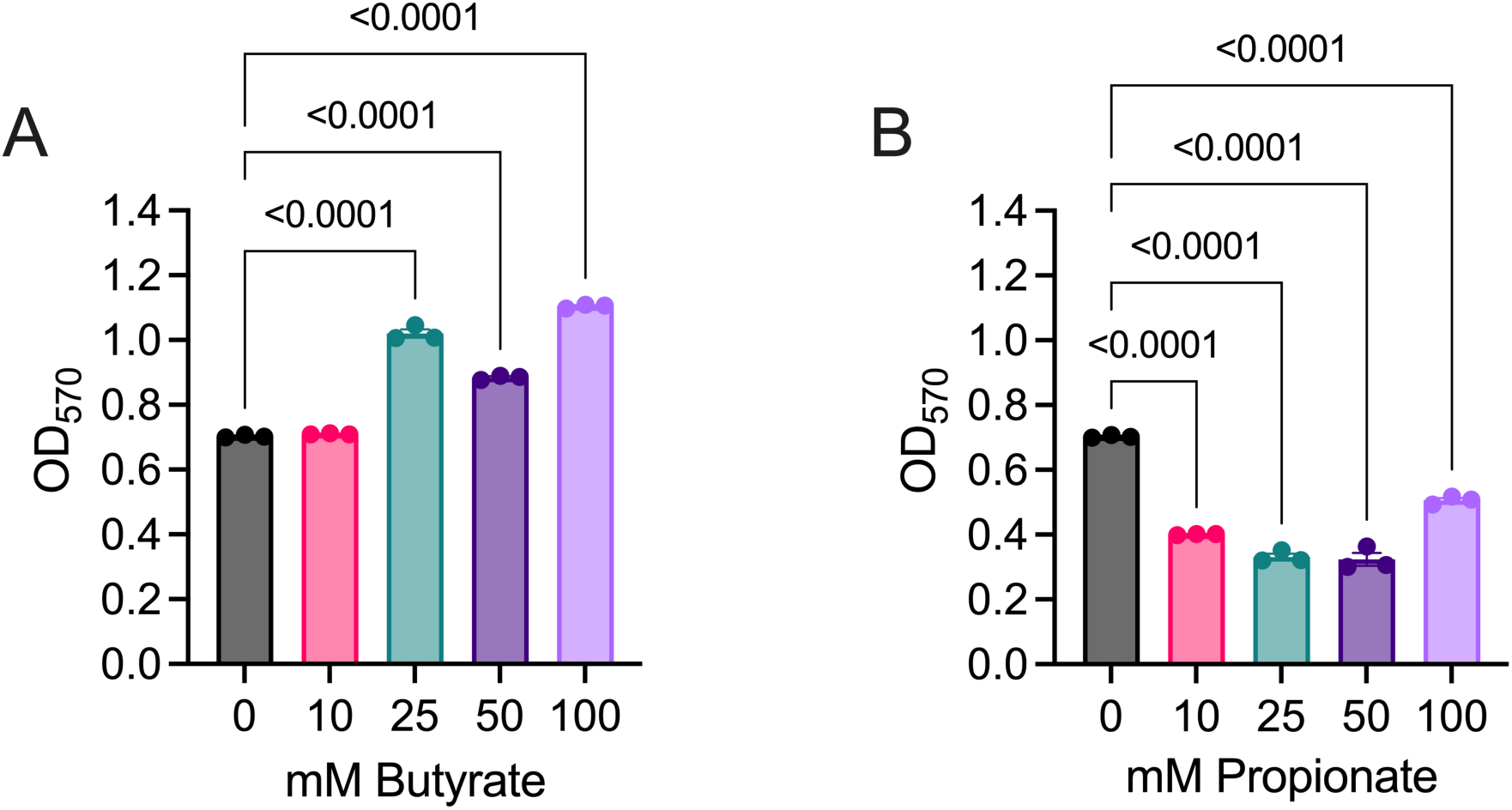
48h anaerobic *S. aureus* biofilms in increasing doses of SCFAs. *S. aureus* JE2 biofilm formation after 48h incubation in LB medium supplemented with indicated concentrations of **(A)** butyrate or **(B)** propionate, as measured by crystal violet staining. Data represent the mean ± standard error of the mean of three independent biological replicates. P-values are from an ordinary one-way ANOVA with Dunnett’s correction for multiple comparisons, with a single pooled variance. All conditions were compared to the 0 mM condition.

**Supplemental Figure 2.**
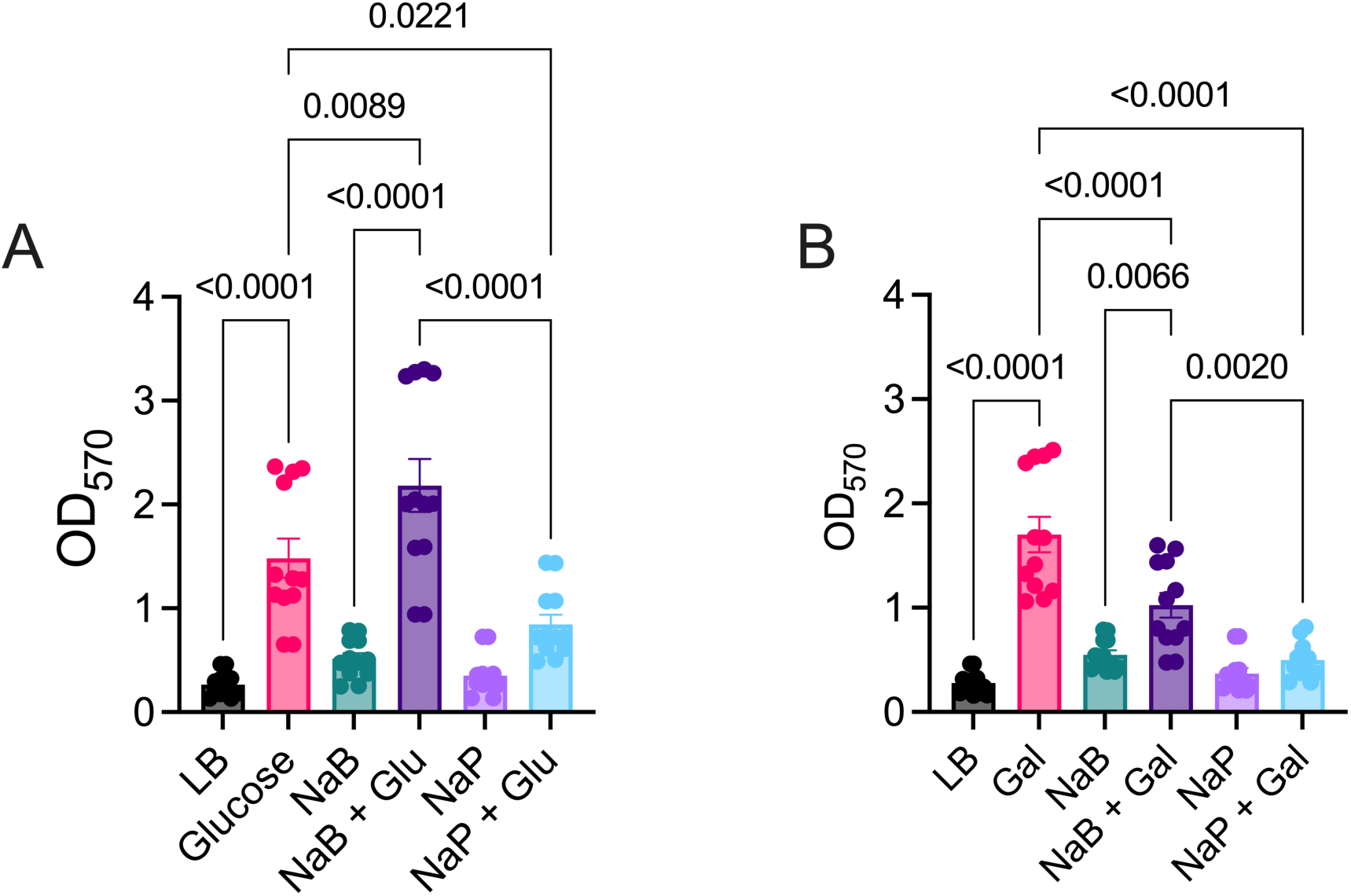
48h anaerobic biofilms of *S. aureus* in 50 mM SCFAs with and without 0.5% w/v sugars. *S. aureus* JE2 biofilm formation after 48h incubation in LB medium supplemented with 50mM SCFAs in the absence or presence of **(A)** 0.5% w/v glucose **(B)** 0.5% w/v galactose, as measured by crystal violet staining. Data represent the mean ± standard error of the mean of twelve independent biological replicates. P-values are from an ordinary one-way ANOVA with Tukey’s correction for multiple comparisons, with a single pooled variance. All conditions were compared to each other, though only relevant comparisons are shown for clarity.

**Supplemental Figure 3.**
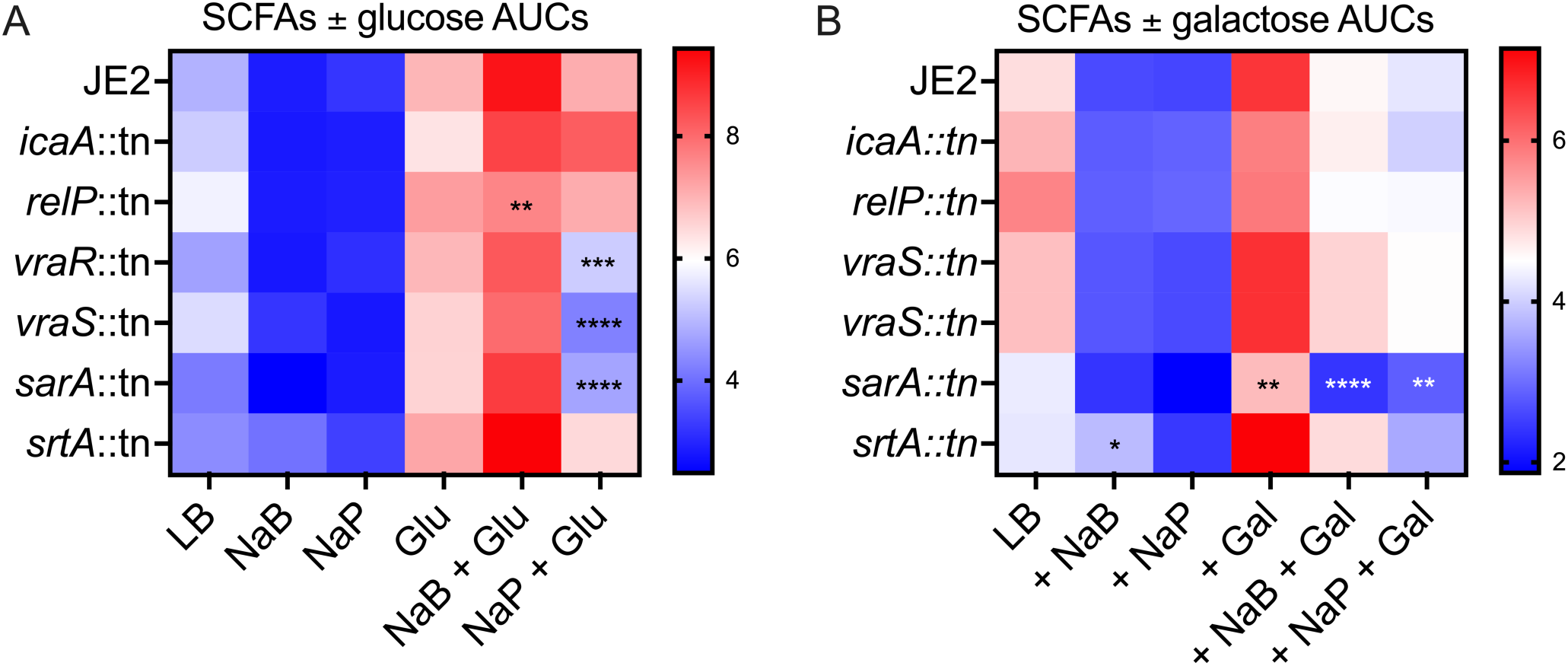
Differential effects of glucose and galactose on SCFA-mediated growth inhibition across *Staphylococcus aureus* strains. Heatmaps of growth curve AUC of *S. aureus* JE2 and the indicated transposon mutants cultured in LB medium with or without 50 mM SCFAs in the absence or presence of **(A)** 0.5% w/v glucose or **(B)** 0.5% w/v galactose. Columns denote media conditions and rows denote strains. Blue indicates reduced growth and red indicates enhanced growth. Data represent the mean ± standard error of the mean of at least three independent biological replicates. P-values are from an ordinary two-way ANOVA with Dunnett’s correction for multiple comparisons, with a single pooled variance. All mutants were compared to JE2 within each media condition. NS (no asterisk) P > 0.05, * P < 0.05, ** P < 0.01, *** P < 0.001, **** P < 0.0001.

**Supplemental Figure 4.**
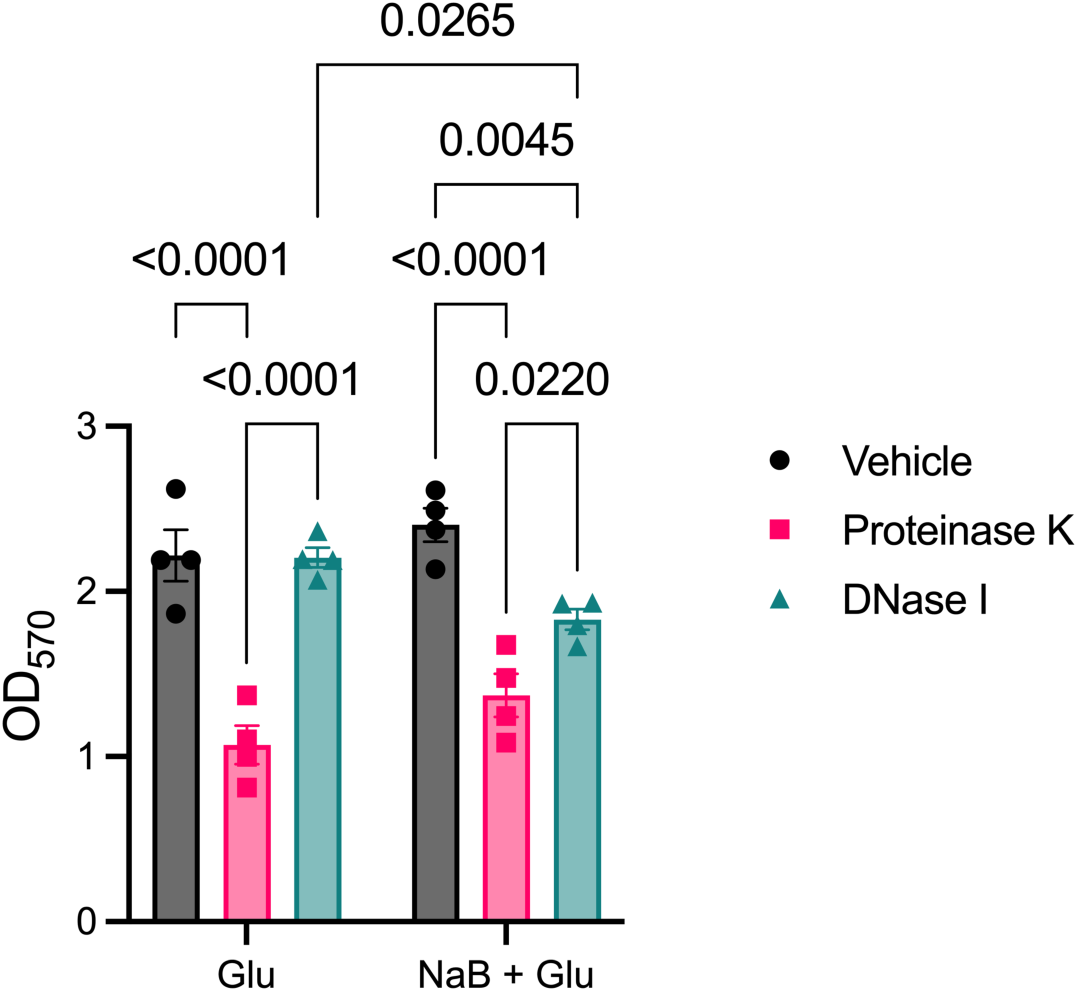
Proteinase K and DNase I reveal distinct contributions of matrix components to biofilm integrity. *S. aureus* JE2 biofilms were established after 24h incubation in LB medium with 0.5% w/v glucose with and without 50mM butyrate. The biofilms were treated with Proteinase K (0.1mg/mL) or DNase I (140U/mL) for 16h at 37°C. Following treatment and washes, the remaining biofilm biomass was quantified via crystal violet staining. Data represent the mean ± standard error of the mean of three independent biological replicates. P-values are from an ordinary two-way ANOVA test with Tukey’s correction for multiple comparisons, with a single pooled variance.

**Supplemental Figure 5.**
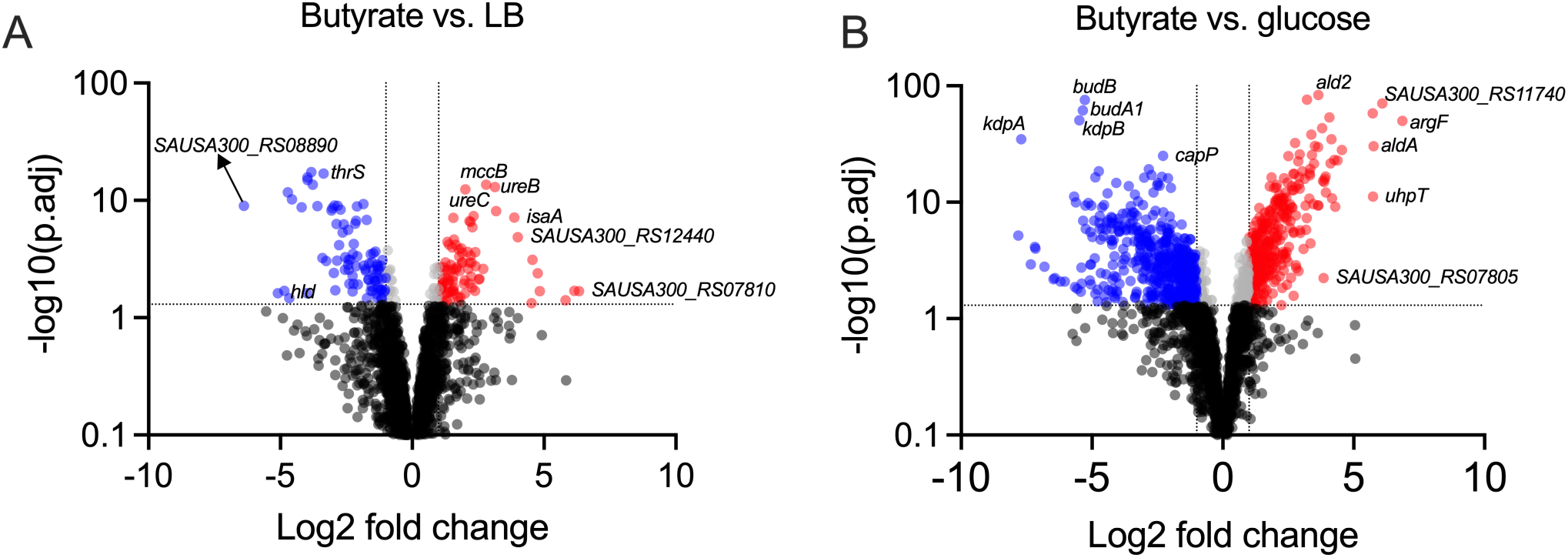
Differential expression analysis of *S. aureus* genes across various media conditions. **(A)** Volcano plot comparing *S. aureus* JE2 gene expression in butyrate versus LB. **(B)** Volcano plot comparing *S. aureus* JE2 gene expression in butyrate versus glucose. Differentially expressed genes are indicated by a minimum of two-fold change in expression at an adjusted p-value < 0.05, with significantly upregulated transcripts colored red and significantly downregulated transcripts colored blue.

**Supplemental Figure 6.**
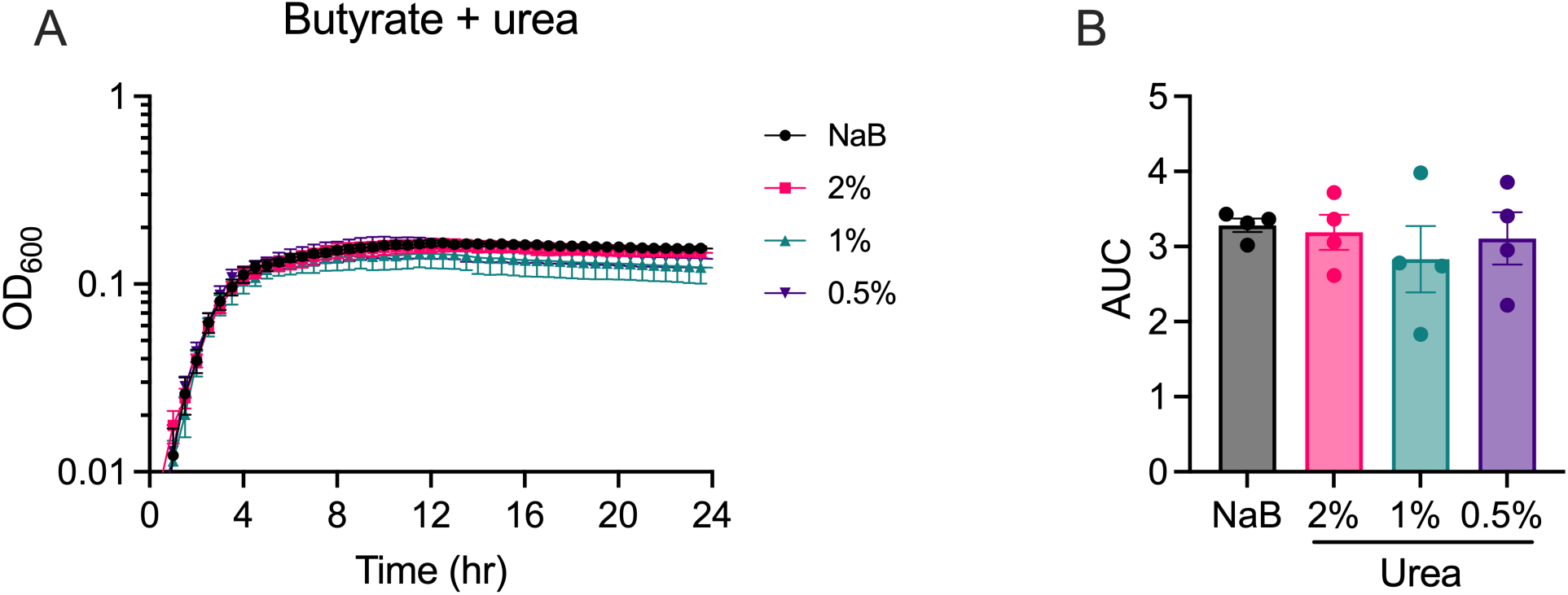
Urea does not rescue *S. aureus* growth when butyrate is present. **A)** Growth curves of JE2 in LB medium with 50 mM butyrate with and without the indicated concentrations of urea. **B)** Area under the curve (AUC) from the growth curves shown in A. Data represent the mean and error bars are the standard error of the mean of four independent biological replicates. All conditions were compared to butyrate and none were significant by ordinary one-way ANOVA with Dunnett’s correction for multiple comparisons, with a single pooled variance.

